# Crosstalk between salicylic acid signaling and the circadian clock promotes an effective immune response

**DOI:** 10.1101/2023.11.21.568095

**Authors:** Olivia Fraser, Samantha J Cargill, Steven H Spoel, Gerben van Ooijen

## Abstract

The rotation of Earth creates a cycle of day and night, leading to predictable changes in environmental conditions. The circadian clock synchronizes an organism with these environmental changes and alters their physiology in anticipation. Prediction of the probable timing of pathogen infection enables plants to prime their immune system without wasting resources or sacrificing growth. Here, we explore the relationship between the immune hormone salicylic acid (SA), and the circadian clock in Arabidopsis. We found that SA altered circadian rhythmicity through the SA receptor and master transcriptional coactivator, NPR1. Reciprocally, the circadian clock gates SA-induced expression of NPR1-dependent immune genes. Furthermore, the clock gene CCA1 is essential for SA-induced immunity to the major bacterial plant pathogen *Pseudomonas syringae*. These results reveal new interactions between the circadian clock and SA signaling which produce an effective immune response. Understanding how and why the immune response in plants is linked to the circadian clock is crucial in working towards improved crop productivity.

## Introduction

The rotation of our planet creates 24-hour cycles in our environments, such as in light levels and temperature. To anticipate these rhythms, the circadian clock has evolved as an internal timekeeping mechanism in almost all eukaryotes. It synchronizes an organism’s behavioral and physiological processes with the rhythmic changes in its surroundings (Dunlap 1999; Reppert and Weaver 2002). Detection of environmental cues feeds into the circadian clock, which then regulates outputs such as rhythmic gene expression. The temporal separation or coordination of process that results from this timekeeping enables the optimization of resource use in an organism’s everyday function (Hodge et al. 2015; Kim et al. 2017). For example, in humans, the circadian clock plays roles in the cardiovascular and nervous systems, gut microbiome, cancer and aging (Bechtold et al. 2010).

As sessile organisms, plants cannot escape unfavorable conditions, evade herbivores, or move to find more distant resources. Therefore, the main challenge for plants is the effective use of available resources to improve fitness. Plants utilize the circadian clock to track and anticipate periodic environmental changes and stressors. This results in the prioritizing of distinct biological responses at different times of day or year according to the anticipated environmental conditions (Dodd et al. 2005; Covington and Harmer 2007; Kim et al. 2017; Creux and Harmer 2019).

In Arabidopsis, circadian rhythms involve an interconnected network of transcriptional/translational feedback loops, in which circadian components activate or repress each other’s activity in a cyclical manner (Creux and Harmer 2019). Expression of the MYB-related transcription factors Circadian Clock Associated 1 (CCA1) and Late elongated Hypocotyl (LHY) peak at dawn. This is followed by the sequential expression of Pseudo-Response Regulator 9 (PRR9), PRR7 and PRR5 throughout the day. The transcription factor, Timing Of Cab expression 1 (TOC1), peaks in the evening, followed by expression of the evening complex (EC), consisting of Early Flowering 3 (ELF3), ELF4 and LUX (Li et al. 2011; Kamioka et al. 2016; Huang and Nusinow 2016). The rhythmic expression of these core clock components in turn regulates many downstream pathways through periodic gene expression. At least 30% of the *Arabidopsis thaliana* (Arabidopsis) transcriptome is under circadian control (Covington et al. 2008). This includes genes involved in regulating flowering, biomass, photosynthesis, water use, temperature stress responses and pathogen defenses (Mizoguchi et al. 2005; Kim et al. 2017; Paajanen et al. 2021).

One of the downstream pathways that the clock modulates is the immune system (Lu et al. 2017). Plant pathogens and pests cause devastating losses to food crops worldwide (Savary et al. 2019). Understanding the mechanisms that underpin plant immunity is critical to improving crop productivity and to meet the exponentially increasing food demand. Pathogen recognition is conferred through two immune pathways: PAMP triggered immunity (PTI) and effector triggered immunity (ETI) (Jones and Dangl 2006; Spoel and Dong 2012). Pathogen recognition by these interconnected pathways leads to accumulation of the immune hormone salicylic acid (SA) that activates the expression of thousands of immune genes, many via the activation of the master transcriptional coactivator, NPR1 (Cao et al. 1994; Cao et al. 1997). NPR1 is responsible for the regulation of pathogenesis related (PR) genes including PR1. In addition to activating local immune responses, SA and NPR1 are also required for the onset of systemic acquired resistance (SAR), which provides long-lasting protection throughout the entire plant against a broad spectrum of pathogens (Malamy et al. 1990; Gaffney et al. 1993; Park et al. 2007; Chanda et al. 2011; Fu and Dong 2013).

Basal levels of SA are under circadian control with SA abundance peaking in the middle of the night, which is thought to prime the immune response for dawn, when plants are most vulnerable to pathogen infection (Melotto et al. 2006; Goodspeed et al. 2012; Zheng et al. 2015). In accordance, Arabidopsis displays enhanced resistance in the subjective morning and increased susceptibility at subjective midnight (Bhardwaj et al. 2011). This disparate resistance is lost in plants overexpressing *CCA1* (*CCA1ox*) and in *elf3-1* mutants, both of which are arrhythmic at the gene expression levels. Furthermore, key immune genes are under circadian regulation, including the pathogen recognition receptors *FLS2* and *EFR*, as well as components of the SA biosynthesis, *ICS1* and *EDS1* (Bhardwaj et al. 2011). Interestingly, rhythmic levels of the active NPR1 monomer have also been observed (Zhou et al. 2015).

There is also evidence for reciprocal modulation of the circadian clock by the immune system, however the nature of clock alteration and the extent to which it is modified are unclear. Arabidopsis infected with *Pseudomonas* or treated with the bacterial elicitor flg22 display a significantly shortened circadian period (Zhang et al. 2013). Moreover, SA accumulation reinforces the circadian clock in an NPR1-dependent manner (Zhou et al. 2015). On the other hand, SA treatment has also been reported to cause a significant reduction of amplitude and delay in circadian phase (Li et al. 2018).

Taken together, these findings point to extensive crosstalk between the immune system and the circadian clock. However, the exact mechanisms and extent of cross modulation are poorly understood.

In this study, we further explore the relationship between SA-dependent immune signaling and the circadian clock in *Arabidopsis*. We found that SA treatment induced significant period shortening of circadian clock gene expression in an NPR1-dependent manner. Conversely, we found that SA-induced expression of *PR1*, a major NPR1 target gene with antimicrobial functions, was higher during the subjective night. This gating of *PR1* induction to the subjective night was lost in clock mutants. Furthermore, clock mutants were unable to establish SA-induced resistance when infected with the bacterial leaf pathogen *Pseudomonas syringae.* Overall, our study reveals novel interactions between SA-dependent immune signaling and the circadian clock, significantly, the importance of NPR1 in clock gene rhythm modulation and the critical role for the clock gene CCA1 in launching an effective immune response.

## Results

### SA treatment shortens the period of the circadian clock

Conflicting reports exist on the effects of SA on the circadian clock in plants. Therefore, we first aimed to consolidate the effect of SA on the clock within our experimental systems and conditions. Leaf discs of plants expressing firefly luciferase (LUC) driven by the rhythmic promoters of either *CCA1* or *TOC1* (*CCA1pro:LUC* and *TOC1pro:LUC,* respectively) were subjected to a range of SA concentrations.

Treatments with 100 µM SA (Figure S1) or 1 mM SA (Figure 1) shortened the periods of both *CCA1* and *TOC1* under constant light conditions. Higher concentrations of SA abolished rhythms while lower concentrations did not significantly alter rhythms (Figure S1).

**Figure 1.**
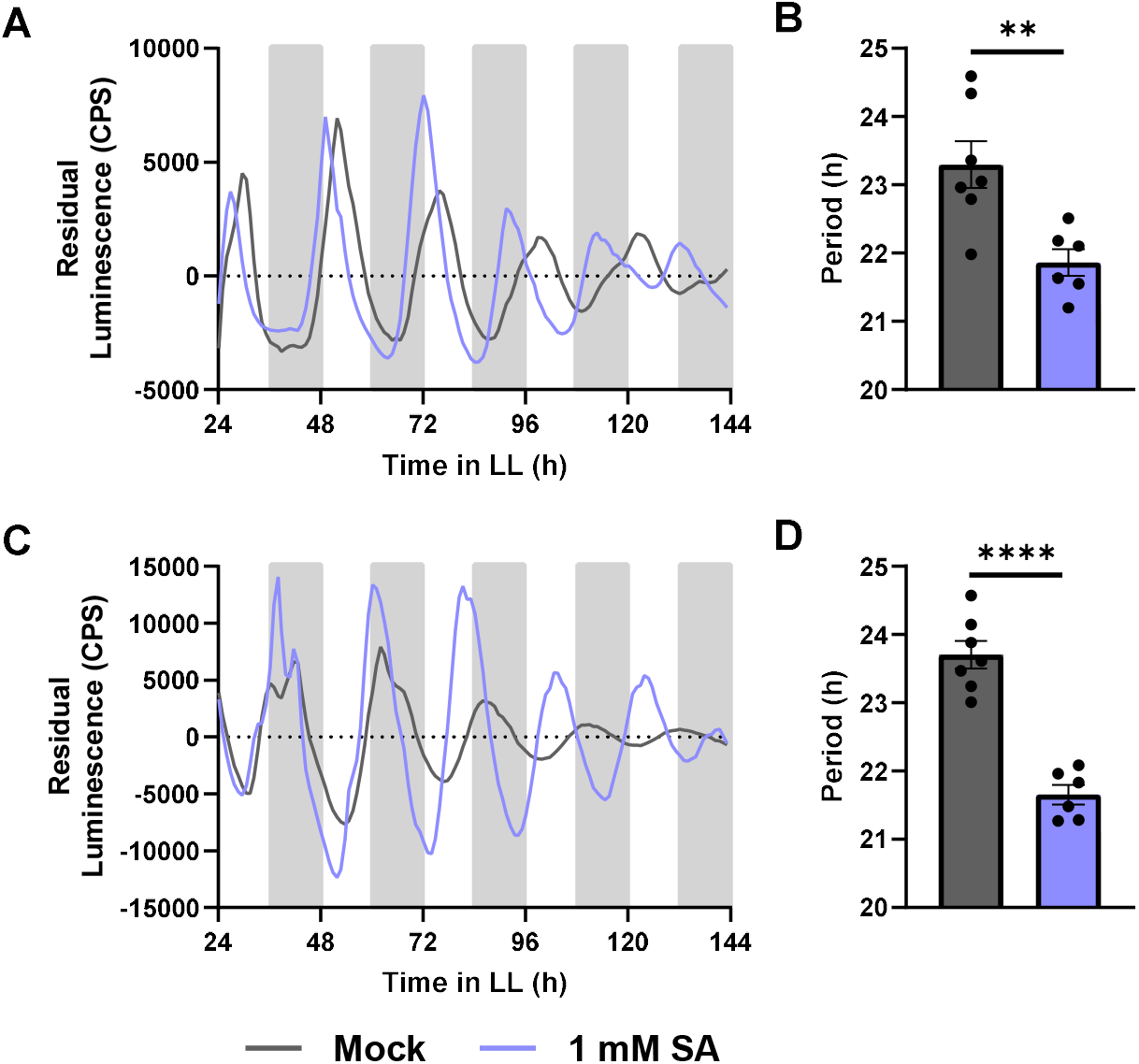
Continuous SA treatment shortens the period of the circadian clock. Promoter activity of CCA1 (A-B) and TOC1 (C-D) was observed by measuring luminescence in CCA1pro:LUC and TOC1pro:LUC leaf disks, respectively. Leaf disks were treated with 1 mM SA (blue) or mock treated with water (grey). The mean period (B and D) was calculated from the 24 – 120 hr time window. An unpaired t-test was performed between each mock and treated data set for CCA1 and TOC1: *p <0.05, **p <0.01, ***p <0.001, ****p<0.0001, unpaired t-test. Error bars indicate mean ± SEM (n = 7). Data are from a single experiment representative of 3 independent repeats.

We next investigated the significance of the timing of SA treatment on circadian rhythms. The bioluminescence of *CCA1pro:LUC* leaf disks was measured in response to treatment with SA at five different time points spanning across a period of 24 hours.

SA treatment significantly shortened the period when applied at time points LL24, LL29, LL34 and LL39 (Figure 2A-D respectively, summarized in Figure 2G). The period shortening was most significant between LL29 and LL34. These results indicate a time-dependency of SA-induced period shortening.

**Figure 2.**
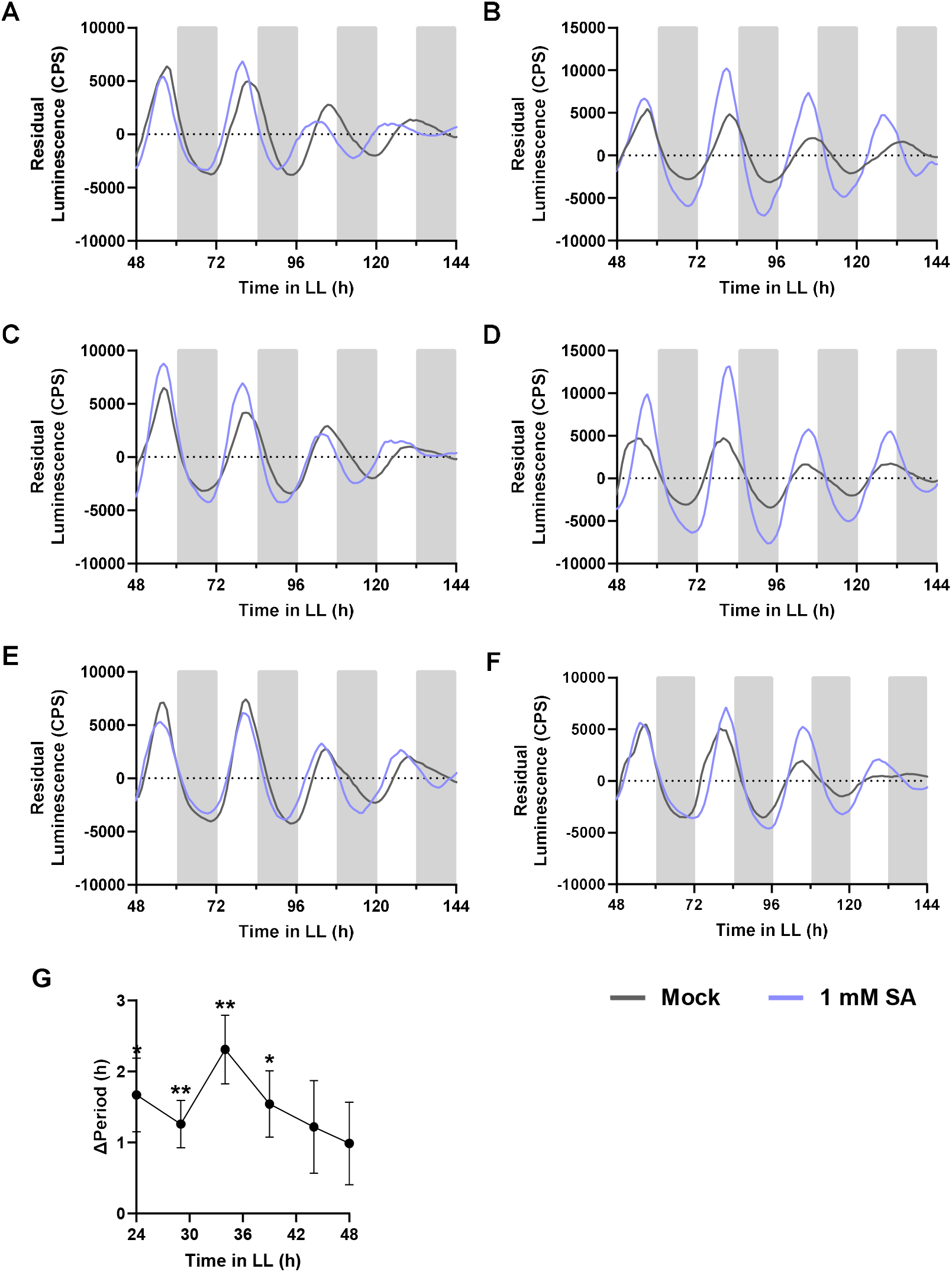
Continuous SA treatment shortens the period of the circadian clock when applied at different times over 24 hours. Promoter activity of *CCA1* (A-F) was observed by measuring luminescence in CCA1pro:LUC leaf disks. Leaf disks were treated with 1 mM SA (blue) or mock (H_2_O) (grey) at 5 different time points over 24 hours: LL24 (A), LL29 (B), LL34 (C), LL39 (D), LL44 (E), LL48 (F). The mean period was calculated from the 48 – 144 hr time window and unpaired t-tests were performed between each mock and treated data set for each time point: *p <0.05, **p <0.01, ***p <0.001, ****p<0.0001, unpaired t-test. Mean ΔPeriod was calculated from the mock and treated data sets and displayed with the results of the unpaired t-test for each time point (G). Error bars indicate mean ± SEM (n = 8). Data are from a single experiment.

Our experiments and those reported previously investigated the effect of continuous exposure to SA on the circadian clock. As basal SA levels are rhythmic (Goodspeed et al. 2012) and pathogen-induced SA accumulation is transient (Ederli et al. 2011), we investigated the effect of transient SA treatment on *CCA1* rhythms. *CCA1pro:LUC* leaf disks were treated with SA at five different time points over 24 hours and the media was replaced with SA-free imaging media 8 hours after treatment. Here, only time points LL34 and LL44 exhibited significant period shortening (Figure 3). This confirms that transient SA treatment also shortens the period of the circadian clock similarly to continuous treatment, though the effect size is somewhat smaller.

**Figure 3.**
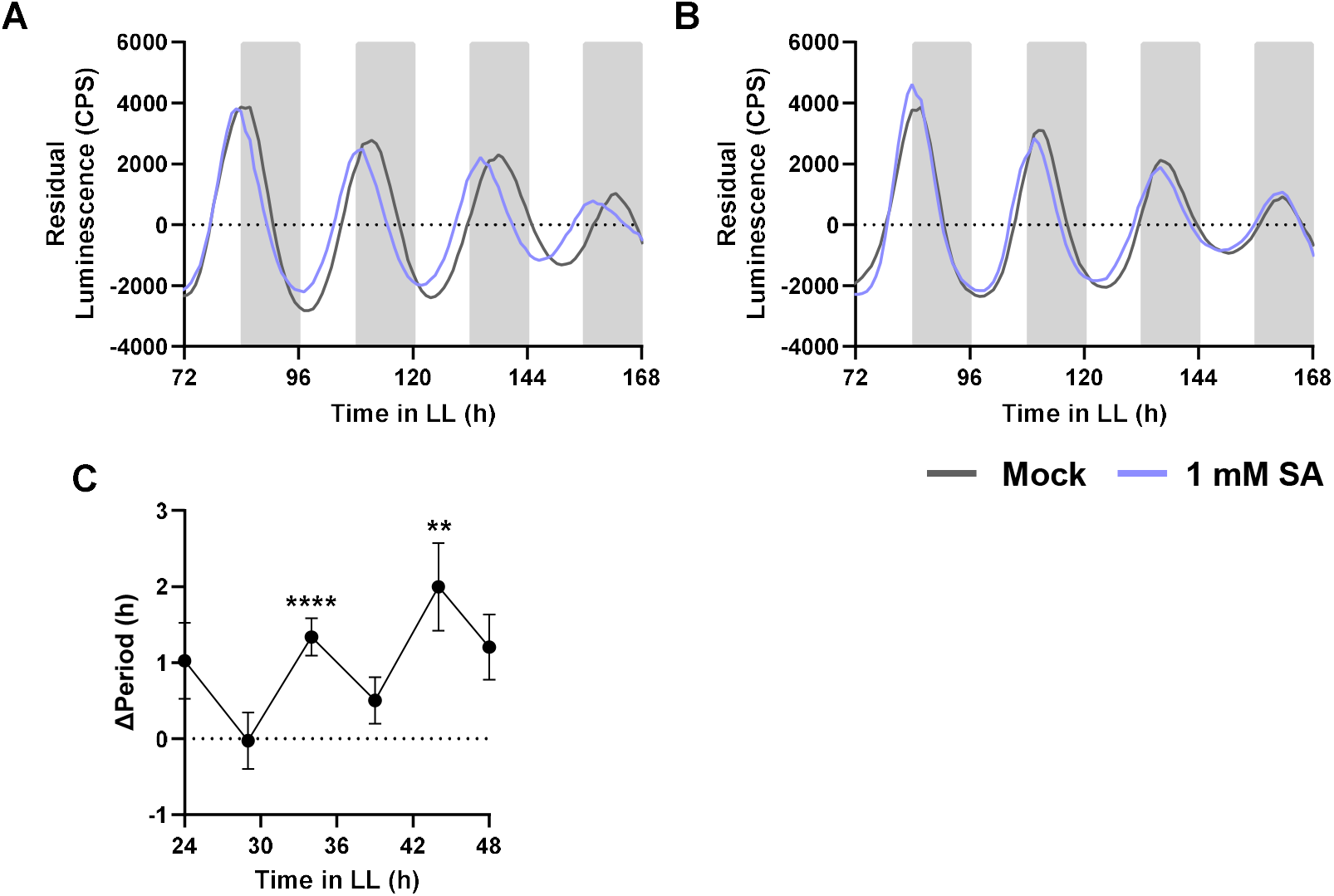
Transient SA treatment shortens the period of the circadian clock when applied at different times over 24 hours. Promoter activity of *CCA1* (A and B) was observed by measuring luminescence in CCA1pro:LUC leaf disks. Leaf disks were treated with 1 mM SA (blue) or mock (H_2_O) (grey) at 5 different time points over 24 hours. The media was removed and replaced with SA-free imaging media 8 hours post treatment. Only luminescence traces of treatment at LL34 (A) and LL39 (B) are shown here to demonstrate significant period shortening and (A) and lack of period shortening (B). The mean period was calculated from the 72 – 168 hr time window and unpaired t-tests were performed between each mock and treated data set for each time point: *p <0.05, **p <0.01, ***p <0.001, ****p<0.0001, unpaired t-test. Mean ΔPeriod (C) was calculated from the mock and treated data sets and displayed with the results of the unpaired t-test for each time point. Error bars indicate mean ± SEM (n = 8). Data are from a single experiment representative of 2 independent repeats.

Collectively, the results in Figures 1-3 suggest that conflicting reports in literature may be due to difference in SA dose dependency and treatment time within experimental setups. Furthermore, they imply that pathogen-induced SA accumulation may modulate the circadian clock in a dose-dependent and time-dependent manner.

### SA-induced period shortening is dependent on NPR1

Our results suggest that fluctuations in SA levels in Arabidopsis could feed back to alter circadian clock rhythms. NPR1 is an SA receptor and master regulator of SA-responsive immune gene expression (Cao et al. 1997). Zhou et al. (2015) previously reported that the effect of SA on *TOC1* is dependent on NPR1. Contrasting results reported by Li et al. (2018) argue that NPR1 antagonizes the clock response to SA treatment. As NPR1 is a regulator of SA-dependent immunity, it may mediate feedback between SA and the clock. However, in previous studies that observed circadian gene expression of NPR1 mutants, no effect on period was found (Zhou et al. 2015; Li et al. 2018).

To investigate this, we observed *CCA1* and *TOC1* rhythms in *npr1-1* mutants. We crossed *CCA1pro:LUC* and *TOC1pro:LUC* with *npr1-1* mutants. First, we compared the rhythms of *CCA1* and *TOC1* in wild-type (WT) versus *npr1-1* mutant background (Figure 4). We found that *npr1-1* mutants had a significantly longer period as observed for both *CCA1* (Figure 4A, B) and *TOC1* (Figure 4C, D). Although the period lengthening effect is small the detailed time resolution provided by luciferase reporters allowed us to evidence a clear effect of NPR1 on circadian gene expression in uninduced conditions

**Figure 4.**
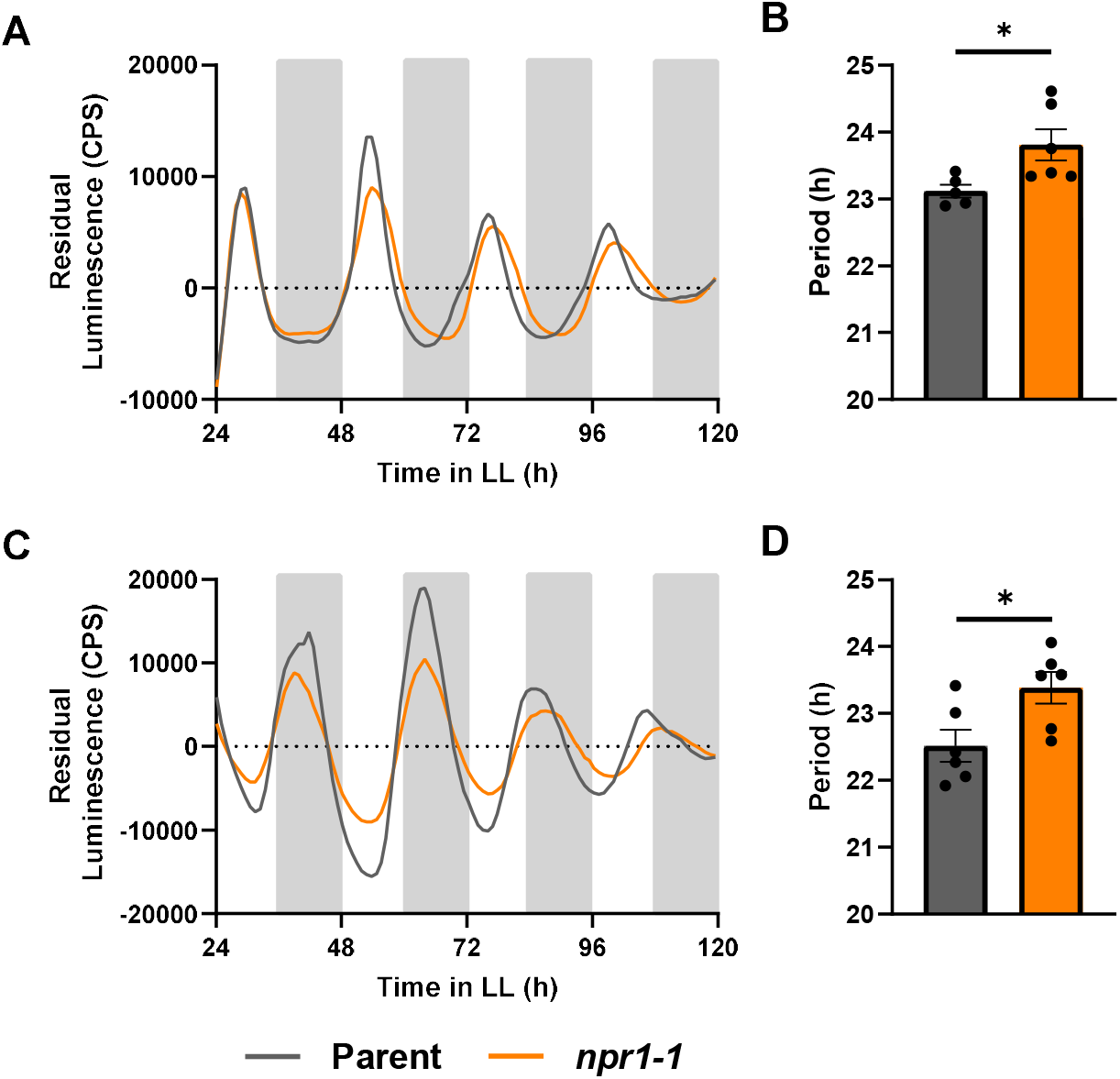
The circadian clock of *npr1-1* displays a longer period than wild type. Promoter activity of *CCA1* (A-B) or *TOC1* (C-D) was measured in the parent line parent line (grey) and the *npr1-1* mutant (orange). *CCA1* promoter activity (A-B) was visualized by measuring luminescence of *CCA1pro:LUC* in wild-type (grey) versus *CCA1pro:LUC* in *npr1-1* (orange) leaf disks. *TOC1* promoter activity (C-D) was visualized by measuring luminescence of *TOC1pro:LUC* in wild-type (grey) versus *TOC1pro:LUC* in *npr1-1* (orange) leaf disks. The mean period was calculated from the 24 – 120 hr time window from traces of *CCA1* (B) and *TOC1* (D). *p <0.05, **p <0.01, ***p <0.001, ****p<0.0001, unpaired t-test. Error bars represent mean ± SEM (n = 7). The data shown here are from a single experiment representative of 3 independent repeats.

To test if NPR1 also mediates the effects of SA on circadian period shortening (Figure 1), we subjected our clock marker lines in the WT and *npr1-1* backgrounds to SA treatment. As opposed to the WT background (Figure1), SA treatment of *CCA1pro:LUC* in *npr1-1* and *TOC1pro:LUC* in *npr1-1* leaf discs did not alter rhythms (Figure 5). Combined, these data demonstrate that NPR1 affects timekeeping and that the effect of SA on the circadian clock is dependent on NPR1. Thus, basal and SA induced NPR1 are part of the input system of the plant circadian clock.

**Figure 5.**
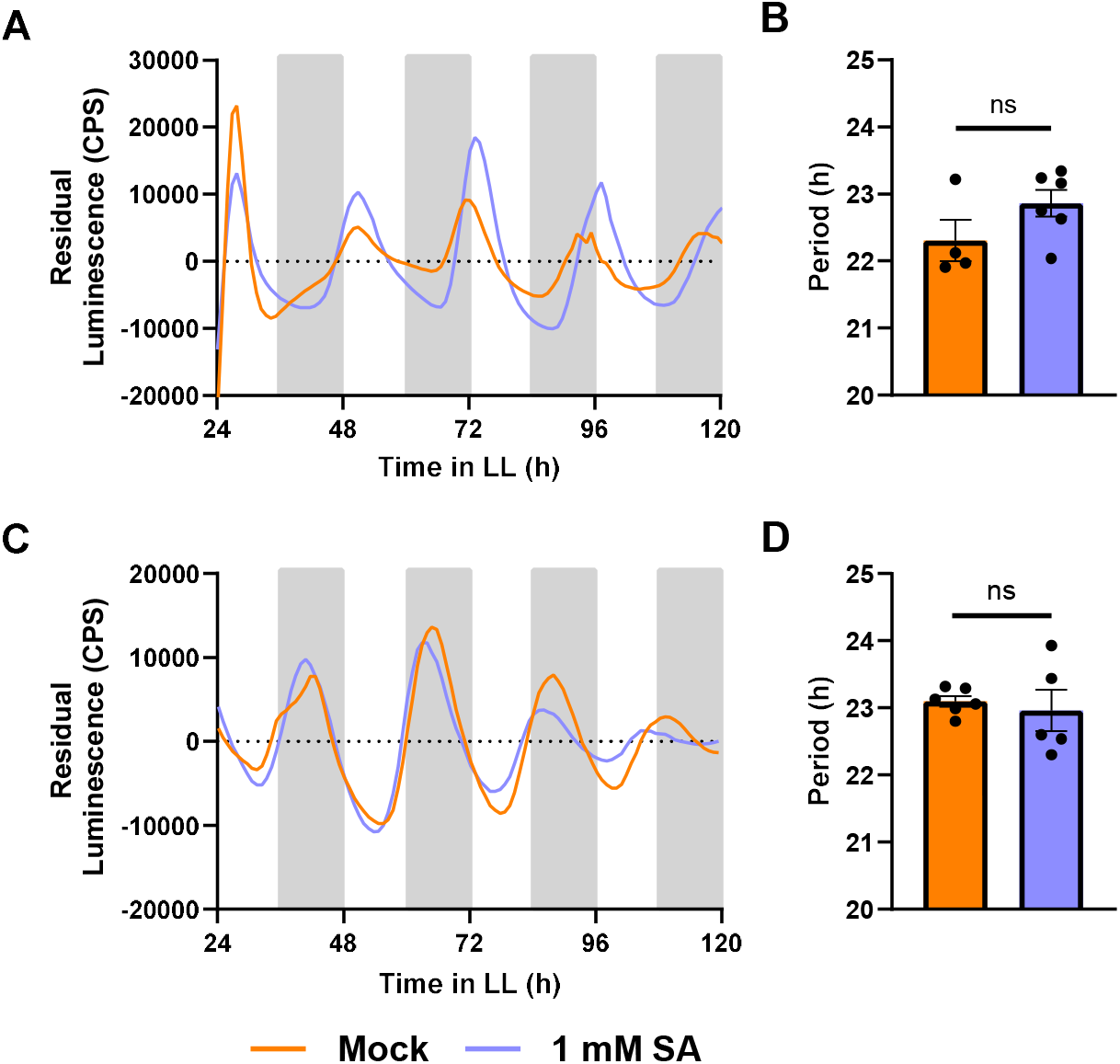
Period shortening by SA is lost in *npr1-1* plants. Promoter activity of *CCA1* (A-B) or *TOC1* (C-D) was observed by measuring luminescence in *CCA1pro:LUC* in *npr1-1* and *TOC1pro:LUC* in *npr1-1* leaf disks, respectively. Leaf disks were treated with 1 mM SA (blue) or mock treated with water (orange). The mean period was calculated from the 24 – 120 hr time window from traces of *CCA1*(B) or *TOC1* (D). *p <0.05, **p <0.01, ***p <0.001, ****p<0.0001 unpaired t-test. Error bars indicate mean ± SEM (n = 7). The data shown here are from a single experiment representative of 2 independent repeats.

### SA-induced PR1 gene expression is gated by the circadian clock

Thus far, NPR1 appears to be one of the key elements in crosstalk between SA levels and the circadian clock. NPR1 is a master regulator of SA-responsive gene expression, including the antimicrobial gene *Pathogenesis-Related 1* (PR1). As endogenous basal levels of SA as well as the abundance of monomeric NPR1 protein exhibit circadian oscillations (Zheng et al. 2015; Zhou et al. 2015), we investigated if *PR1* gene expression is also rhythmic. We measured PR1 expression in wild-type plants by sampling every 3 hours for two days. In the absence of an immune inducer, basal *PR1* gene expression remained very low and did not exhibit rhythmicity (Figure S2). However, when we subjected plants to SA at 3-hour intervals, we found that SA-induced *PR1* expression was consistently higher during the subjective night than the subjective day (Figure 6). Thus, the circadian clock might gate SA-induced *PR1* expression.

**Figure 6.**
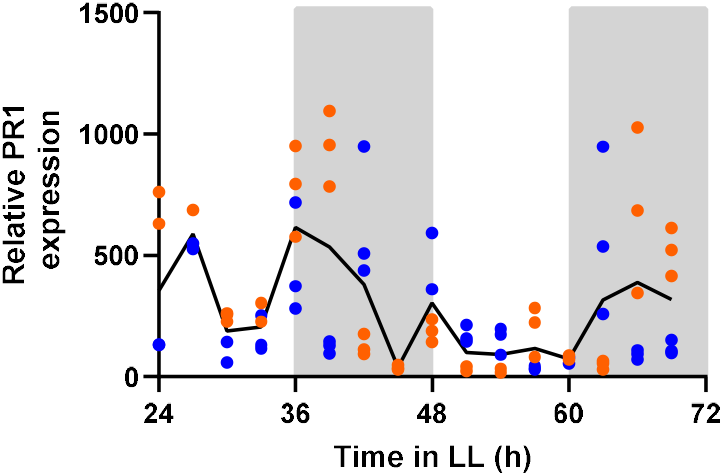
SA induced PR1 expression is higher during the subjective night. Col-*0* seedlings were grown on MS plates under 12/12 light/dark conditions for 14 days before being released into constant light (LL). Every 3 hours, plates were sprayed with 1 mM SA. Relative expression of PR1 was measured 6 hours post treatment. The data points are technical replicates of quantitative PCR results combined from two biological replicates (orange versus blue data points; n = 3 each). The line represents mean values of all 6 replicates.

To investigate this further, we compared SA-induced *PR1* expression levels at subjective dawn versus dusk in wild-type as well as in arrhythmic *CCA1ox* plants (Wang and Tobin 1998) and *cca1lhy* mutants that exhibit a shortened period (Zhang et al. 2013). As expected, wild-type plants exhibited stronger induction of *PR1* gene expression in response to SA treatment at dusk (Figure 7A). Higher *PR1* expression at dusk compared to dawn was reinforced in *CCA1ox* plants (Figure 7B), but completely lost in the *cca1lhy* mutant (Figure 7C). These data indicate that core clock component CCA1 gates SA-dependent gene expression and enhances responsiveness to SA at dusk.

**Figure 7.**
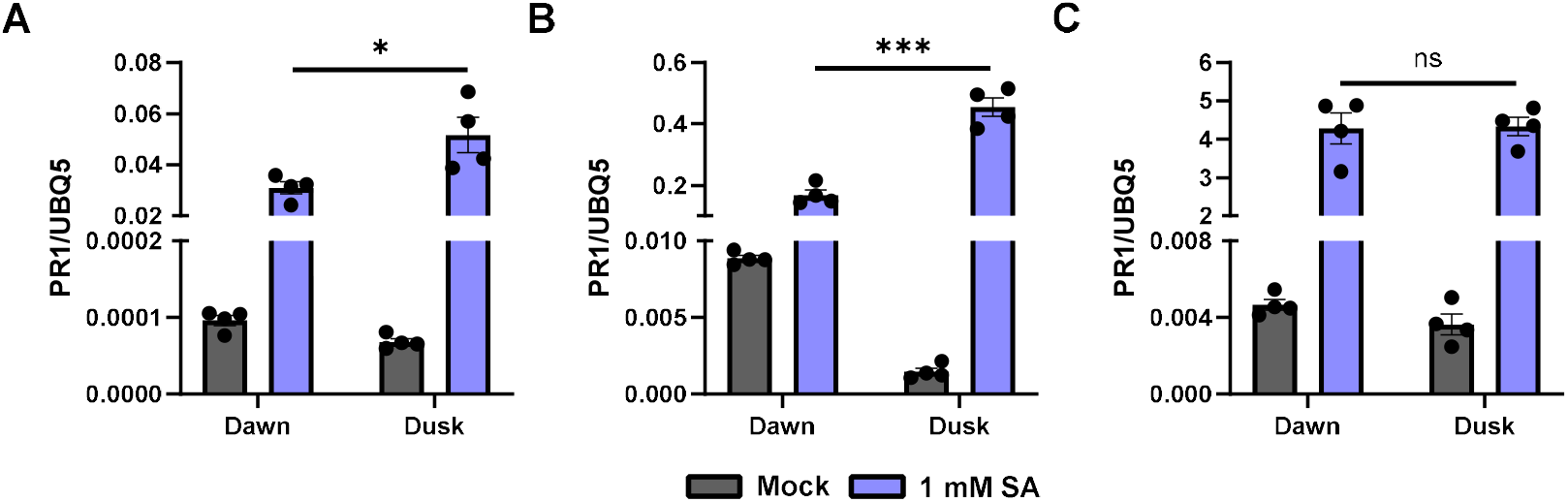
CCA1 expression is required for the disparate expression of PR1 in response to SA treatment. Col-*0* (A), *CCA1ox* (B) and *cca1lhy* (C) seedlings were sprayed with 1 mM SA or mock (H_2_O) at subjective dawn versus dusk. Seedlings were sampled 6 hours after treatment and gene expression was measured. *p <0.05, **p <0.01, ***p <0.001, ****p<0.0001, unpaired t-test. Error bars indicate mean ± SEM (n = 4). The data shown here are from a single experiment, representative of 2 independent experiments.

### CCA1 is required for SA induced resistance

Our results to this point demonstrate evidence of crosstalk between SA signaling and the circadian clock. In spite of the evidence for clock regulation of SA levels and immune gene expression (Bhardwaj et al. 2011; Goodspeed et al. 2012; Zheng et al. 2015), no direct evidence exists for circadian involvement in the process of SA-induced resistance to pathogens. Therefore, we investigated the role of the circadian clock in establishing SA-induced resistance. We sprayed wild-type (Col-0), *CCA1ox*, *cca1lhy*, *TOC1ox* (arrhythmic: Makino et al. 2002) and *toc1-101* (short period: Kikis et al. 2005) plants with SA at subjective dawn to mimic pathogen infection, followed 24 hours later by infiltration with *Pseudomonas syringae pv. maculicola (Psm)* ES4326. As expected, wild-type plants displayed a significantly lower level of infection after treatment with SA (Figure 8). *TOC1ox* plants and *toc1-101* mutants also displayed SA-induced resistance, indicating that neither TOC1 expression nor its rhythmicity is required for SA-mediated resistance. However, SA-induced resistance was not observed in *CCA1ox* plants or in *cca1lhy* mutants. These results indicate that both *CCA1* expression as well as its rhythmic expression are required to establish SA-dependent disease resistance. These results further evidence the critical role for CCA1 in linking the circadian clock, SA signaling, and immunity.

**Figure 8.**
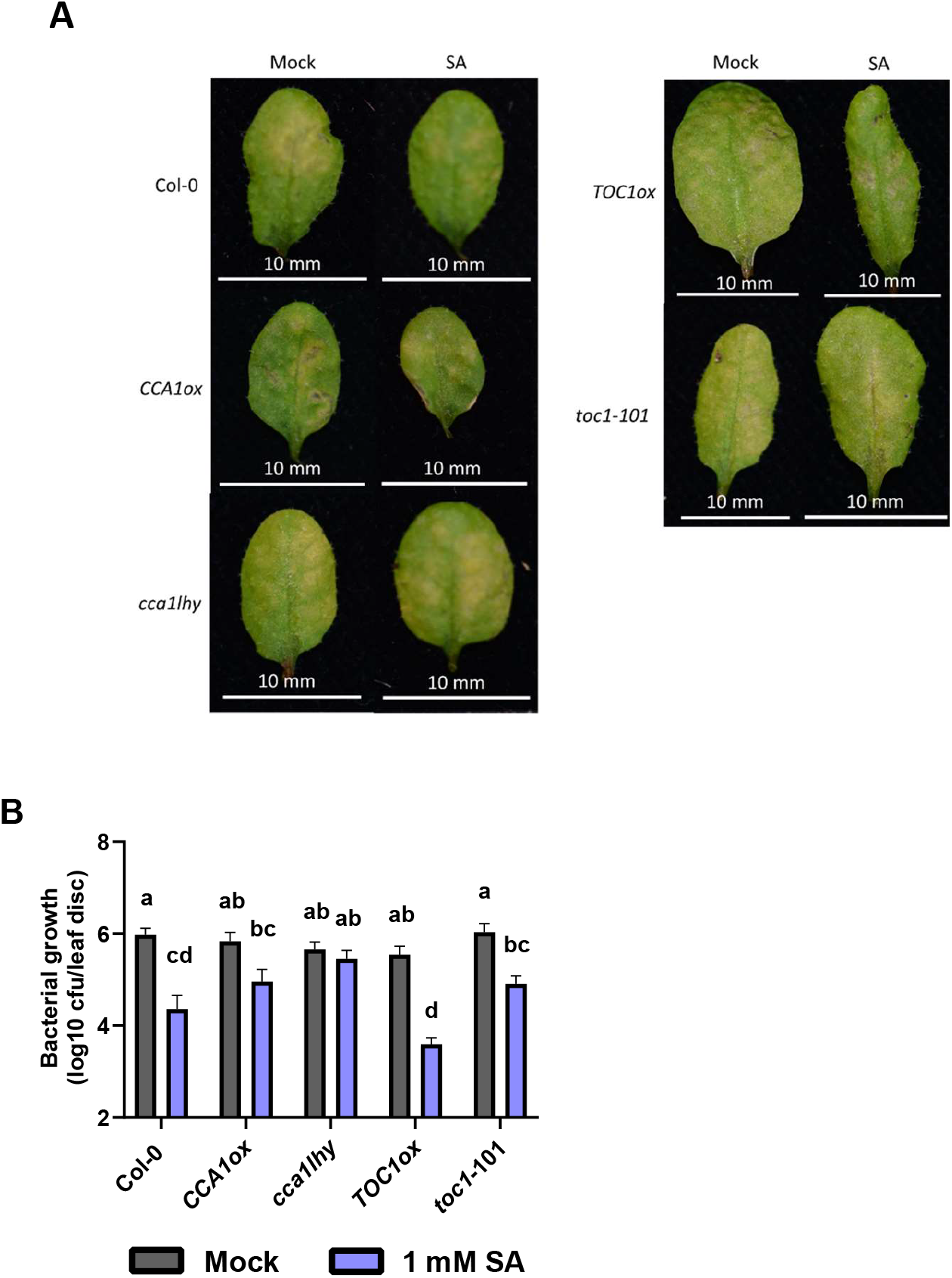
CCA1 circadian mutants do not display SA induced resistance to *Pseudomonas.* Col-*0*, *CCA1ox, ccalhy*, *TOC1ox* and *toc1*-101 plants were grown under 12/12 light/dark conditions for 3 weeks. At dawn on day 21 (LL0), the plants were released into constant light conditions. At LL24 the plants were sprayed with 1 mM SA. At LL48 the plants were infected with *Pseudomonas.* The level of bacterial growth was measured 3 days post infection. (A) Leaves representative of the visual median level of infection at 3 days post infection for each genotype and treatment. (B) The infection level at 3 days post infection expressed by colony forming units (CFU, y-axis is logarithmic) in leaf disk extracts. Error bars represent mean ± SEM (n = 8 individual leaves). A two-way ANOVA and Tukey’s HSD Test were preformed to analyse the differences in bacterial growth between treatments and genotypes. The letters indicate significant differences between groups (p < 0.05). The data shown here are from a single experiment, representative of 3 independent experiments (Col-*0* versus *CCA1ox* or *cca1lhy*) or two independent experiments (Col-*0* versus *TOC1ox* or *toc1-101*).

## Discussion

Plant pathogens are responsible for large reductions in agricultural productivity (Savary et al., 2019), A functioning circadian clock has been shown to enhance resistance to pests and pathogens both pre- and post-harvest (Goodspeed et al. 2012, 2013). The regulation of the immune system by the circadian clock has long been theorized to prime the immune system to the time of day that pathogen attack is most likely, thereby conserving the plants resources (Goodspeed et al. 2012). The alteration of circadian rhythms during infection may function to prioritize resource allocation towards an immune response by resetting the circadian clock to a time of day that facilitates this through its regulation of other plant processes.

The results presented in this study indicate that the immune hormone SA modulates that circadian clock, inducing NPR1-dependent period shortening. Previous studies have reported conflicting results regarding the effect of SA on circadian clock rhythms. Zhang et al. (2013) found that treating seedlings with the SA mimic benzo(1,2,3)thiadiazole-7-carbothioic acid (BTH) had no effect on *CCA1* promoter rhythms. In a similar study, Zhou et al. (2015) found that treatment with SA reinforced TOC1 rhythms but not CCA1 rhythms in 3-week-old Arabidopsis.

Li et al. (2018) monitored the effect of transient (4 h) SA treatment on clock gene rhythms, observing a significant reduction in amplitude and a delay in phase for CCA1, TOC1 and LHY rhythms. This delay in phase was larger when SA treatment began closer to subjective dawn, though no such pattern was observed in the amplitude reduction. None of these studies observed any effect of long-term SA exposure on period.

To understand the discrepancies of the effect of SA signaling on the clock, we investigated the effect of both continuous and transient SA treatment on clock gene expression using a range of treatment times and consistent methodology. Unlike previous studies, we observed consistent period shortening of *CCA1* and *TOC1* expression upon both transient and continuous treatment with SA. Notably, the length of SA treatment determined the extent to which clock alterations occurred with longer treatment having more pronounced effects. We did not observe a significant effect on phase induced by either continuous or transient SA treatment. We found that the magnitude of the period shortening increased when SA exposure began between LL29 and LL34. These results contrast those reported by Li et al. (2018), who did not observe SA-induced period shortening upon transient treatment but did report a lager phase shift effect around subjective dawn. Therefore, although our study and the previous studies do not agree on the exact effect of SA treatment on clock gene rhythms, we contribute to the conclusion that there is time-dependent feedback between the immune system and the circadian clock via SA signaling.

Li et al. (2018) found that the *npr1-1* mutant displayed a larger reduction in the amplitude of CCA1 and LHY upon SA treatment, suggesting that NPR1 acts as an antagonist of the effects of SA on the clock. Zhou et al. (2015) also observed that NPR1 was essential for the effect of SA on the clock as the *npr1-1* mutant did not exhibit the SA induced reinforcing of TOC1 rhythms. We found that the SA receptor NPR1 was crucial for the SA-induced period shortening. The amplitude effects we observed were not consistent between replicates and experiments, and therefore we chose not to report on them here. Significantly, we also found that the *npr1-1* mutant displayed a longer period for both *CCA1* and *TOC1* expression in uninduced conditions. This novel finding indicates that basal NPR1 levels, as well as SA-induced NPR1, modulates the circadian clock. The NPR1-dependent SA alteration of circadian clock gene rhythms may function to harness that clock’s modulation of other cellular processes in order to prioritize an immune response when pathogen attack is detected.

Existing literature reports the importance of *CCA1* in launching an immune response to primary bacterial infection. Zhang et al. (2013) report that CCA1ox plants exhibit increased susceptibility to Pseudomonas infection compared to wild type. Conversely, it has also been observed that *CCA1* overexpression confers increased resistance to oomycete pathogen *Hyaloperonospora arabidopsidis (Hpa)* while the *cca1* mutant displayed increased susceptibility (Wang et al. 2011; Zhang et al. 2013).

Our study aimed to expand upon this knowledge of CCA1’s role in immunity by investigating the potential involvement of core clock genes in SA-induced immunity. We demonstrate the importance of the circadian clock in launching an effective immune response and expand upon the critical role of *CCA1* in this coordination. We found that SA-induced *PR1* expression is higher during the subjective night and expression of the clock gene *CCA1* is required for the disparate levels of SA-induced *PR1* expression.

In addition, we observed that *CCA1* is also crucial for the downstream function of establishing SA-induced resistance to Pseudomonas. We found that priming with SA did not induce resistance in *CCA1ox* or *cca1lhy* as it did in Col-0, suggesting that *CCA1* is crucial for the resistance response following SA treatment. We did not observe increased susceptibility to Pseudomonas in *CCA1ox* as reported by Zhang et al. (2013). CCA1 is a key morning clock gene and master regulator of circadian rhythms. Plants overexpressing *CCA1* have an arrhythmic clock under LD and LL conditions (Wang and Tobin 1998). *TOC1ox* (also arrhythmic [Makino et al. 2002]) displayed SA induced resistance to Pseudomonas, suggesting that the lack of SA induced resistance in *CCA1ox* is not due to overall clock arrhythmia. As the *cca1lhy* mutant also did not display an SA-induced response, regulated *CCA1* expression levels appear to be important for establishing SA-induced resistance. Overall, these findings indicate that a function of *CCA1* outside of its role as a core clock component is responsible for its effect on the SA-induced immune response.

From our study, it is evident that SA utilizes NPR1 for modulating rhythmicity of core clock components. NPR1 is crucial for many immune processes downstream of SA. The *npr1-1* mutant displays increased susceptibility to infection, a lack of PR gene expression upon infection/SA treatment and an inability to establish resistance following priming with SA (Cao et al. 1994; Cao et al. 1997). As we have shown that vice versa the circadian clock is also involved in regulating SA induced PR gene expression and resistance to Pseudomonas, it would be interesting to investigate whether NPR1 plays a role in this interaction, potentially facilitating crosstalk between the circadian clock and the immune system in both directions. Additionally, as redox regulation of NPR1 conformation and localization is a key step in its activation (Mou et al. 2003; Tada et al. 2008), daily redox rhythms (Edgar et al. 2012; Lai et al. 2012) may control the interplay between the circadian clock and NPR1.

Our study provides evidence of the reciprocal regulation between the circadian clock and SA signaling. We show the importance of NPR1 in this interaction and that clock gene *CCA1* is key for regulation of SA-induced immune responses. The findings presented here further our understanding of the interactions between the circadian clock and the immune system and how they work together to produce an effective immune response. Continued study of the systems controlling plant immunity is essential to prioritize plant health and thus food security.

## Materials and methods

### Plant materials and growth conditions

*Arabidopsis thaliana* were grown on soil for 3-4 weeks, unless otherwise stated, in growth cabinets (Percival Scientific Inc.) set at constant 21ᵒC with a light intensity of 70-100 µmoles m^-2^ s^-1^ (Fusion 18W T8 2ft Triphosphor Fluorescent Tube 4000K) under 12 h light/12 h dark. All wild type and mutant plants were from the Col-0 genetic background and were described previously: *npr1-1* (Cao et al. 1994); *CCA1ox* (Wang and Tobin 1998); *TOC1ox* (Makino et al. 2002); *cca1lhy* (Zhang et al. 2013); *toc1-101* (Kikis et al. 2005), *CCA1pro:LUC* (Gould et al. 2018) and *TOC1pro:LUC* (Voß et al. 2015).

*CCA1pro:LUC npr1-1 and TOC1pro:LUC npr1-1* plants were made by crossing *CCA1pro:LUC* with *npr1-1* and *TOC1pro:LUC* with *npr1-1* respectively. To obtain homozygous lines of *npr1-1* plants, individuals in the F1 and F2 generations were evaluated by polymerase chain reaction (PCR) genotyping and restriction fragment length polymorphism (RFLP) analysis. DNA was extracted using a one-step protocol previously described in Edwards et al. (1991). For PCR amplification, 0.5 μl of the DNA extract was added to 10 μl of 5X GoTaq Reaction Buffer (Promega) and 1 μl of each primer (10 μM, NPR1 F: CTCGAATGTACATAAGGC, NPR1 R: CGGTTCTACCTTCCAAAG [Cao et al. 1997]), to a total volume of 20 μl. PCR conditions were 95°C for 5 minutes, 35 cycles of 95°C for 30 seconds, 55°C for 30 seconds and 72°C for 1 minute, followed by 72°C for 5 minutes. PCR products were digested with 0.4 μl of restriction enzyme NlaIII for 2 hours at 37°C (New England Biolabs). Digested PCR products were run on 2% agarose gels with SYBR Safe (Invitrogen) at 100 V and were imaged using the Odyssey FC imaging system (LI-COR).

Homozygous *npr1-1* lines were tested for homozygosity of the luciferase transgene (either *CCA1pro:LUC* or *TOC1pro:LUC*) in F3. A minimum of 50 seeds of each line were visualized for luminescence by sowing one seed per well, on top of 200 μl 0.5 MS in black 96-well plates (Greiner bio-one, 655075). Seedlings were grown for 10 days in an incubator (Sanyo) under long-day conditions. 20 μl of 1 mM D-luciferin (Biosynth AG) was added to each well and luminescence was visualized using the ALLIGATOR luminescence imaging system by exposing the camera for 8 minutes. Ratios of luminescence were assessed: homozygous lines which displayed luminescence in all seedlings were used for subsequent experiments.

### Bioluminescence assay of leaf discs

*CCA1pro:LUC, TOC1pro:LUC, CCA1pro:LUC npr1-1* and *TOC1pro:LUC npr1-1* were grown on soil under LD conditions (see plant materials and methods). At 3-4 weeks old, leaf disks were taken from leaves 5-6 and placed in the wells of a white, flat-bottom 96-well plates (Greiner bio-one, 655075) with the adaxial side up. Each well also contained 200 µl of filter sterilized imaging solution: 0.5 MS pH 7.5, 50 µg/ml ampicillin, 1.5 mM D-luciferin and SA/H_2_O to the desired final concentration. The plate was sealed with a clear, gas permeable lid (4titude, 4ti-0516/96). Luminescence was measured by a LB942 Tristar2 plate reader (Berthold Technologies Ltd) every 50 minutes for 3 seconds per well. Leaf disks were kept under continuous red (630 nm) and blue (470 nm) LED light at 17.5 µmoles m^-2^ s^-1^ each at 20-21ᵒC. Results were analyzed using GraphPad Prism and Biodare2 and period was estimated using the Fast Fourier Transform-Non-Linear Least Squares (FFT-NLLS) function.

### SA treatment for quantitative PCR in wildtype over 48 hours

Wild type plants (Col-0) were grown on MS agar plates under 12 h light/12 h dark conditions at 21ᵒC with a light intensity of 70-100 µmoles m^-2^ s^-1^ (Fusion 18W T8 2ft Triphosphor Fluorescent Tube 4000K). After 14 days, the seedlings were released into constant light (LL). Plates were removed from the growth cabinet and sprayed with 1 mM SA or mock (H_2_O) at 3-hour intervals beginning at LL24 and finishing at LL72. After spraying, the plates were put into separate growth cabinets from the untreated plates. The seedlings were harvested and frozen in liquid nitrogen 6 hours after treatment.

### SA treatment for quantitative PCR in clock mutants

Wild type (Col-0), *CCA1ox* and *cca1lhy* plants were grown on MS agar plates under 12 h light/12 h dark conditions at 21ᵒC with a light intensity of 70-100 µmoles m^-2^ s^-1^ (Fusion 18W T8 2ft Triphosphor Fluorescent Tube 4000K). Plates were placed under split entrainment conditions in separate growth cabinets, aligning the time of dawn and dusk between the two cabinets. After 14 days, the seedlings were released into LL. Plates were sprayed with 1 mM SA or mock (H2O) at subjective dawn (LL24) or dusk (LL36). The seedlings were harvested and frozen in liquid nitrogen 6 hours after treatment.

### RNA extraction and cDNA synthesis

Seedlings were manually ground to a fine powder in liquid nitrogen, before being homogenized in RNA extraction buffer (100 mM LiCl, 100 mM Tris pH 8, 10 mM EDTA, 1% SDS). An equal volume of phenol/chloroform/isoamyl alcohol (25:24:1) was added, followed by votexing and centrifuging at 13,000 rpm for 5 min. The aqueous phase was transferred to a tube containing an equal volume of chloroform/isoamyl alcohol (24:1), followed by vortexing and centrifuging at 13,000 rpm for 5 min. This step was repeated before the aqueous phase was transferred to a tube containing 1/3 vol of 8 M LiCl and incubated overnight at 4ᵒC. The samples were centrifuged at 13,000 rpm at 4ᵒC for 15 min. The supernatant was removed and the pellet was washed twice in ice-cold (-20ᵒC) 70% ethanol. The pellet was then dissolved in 400 µl H_2_O for 30 min on ice. 40 µl of NaAc (pH 5.3) and 1ml of ice-cold 96% ethanol was added and the solution was incubated for at least 1 h at -20ᵒC. The tubes were centrifuged at 13,000 rpm for 15 min at 4ᵒC and the pellet was washed twice with ice-cold 70% ethanol. The final pellet was dissolved in 50 µl H_2_O. The resulting RNA was quantified using a NanoDrop spectrophotometer and diluted to standardize the concentrations across all samples. SuperScript II reverse transcriptase (Invitrogen) was used to perform reverse transcription according to the manufacturer’s instructions.

### Quantitative PCR

For SA-induced gene expression in wild-type over 48 hours (Figure 6 and S2), cDNA was diluted 20-fold and qPCR was performed in a 5 µl reaction using SYBR Green and gene specific primers on a QuantStudio 5 PCR machine.

For SA-induced gene expression in clock mutants at subjective dawn and dusk (Figure 7), cDNA was diluted 20-fold and qPCR was performed in a 10 µl reaction using SYBR Green and gene specific primers on a StepOne Plus Real Time PCR machine

### Pseudomonas disease assay

Col-0, *CCA1ox, cca1lhy, TOC1ox* and *toc1-101* were grown on soil under LD conditions (see plant materials and methods). At 3.5 weeks old, plants were released into LL. At subjective dawn (LL24), plants were sprayed with 1 mM SA or mock (H_2_O). At LL48, 2 leaves (leaves 4-6) were pressure infiltrated with *Pseudomonas syringae pv. maculicola (Psm)* at OD_600_= 0.005, using a 1 ml syringe. The bacteria were left to infect the plants for 3 days, before leaf disks were harvested from the infiltrated leaves. The leaf disks were homogenized and diluted in 10 mM MgSO_4_, then plated onto LB plates (10 mM MgSO_4_ and 50 µg/ml streptomycin). Plates were incubated for 2 days at 28ᵒC, before calculating colony forming units (CFU) per leaf disk.

## Acknowledgements

We would like to thank the members of the van Ooijen Lab and the Spoel Lab at the University of Edinburgh for their assistance with this study. We would like to thank Karen Halliday for kindly giving us the clock mutant lines and luciferase lines used in this study.

This research was carried out with resources provided by the Edinburgh Plant Growth Facility, a specialist service provider for plant growth at the University of Edinburgh.

## Author contributions

Conceptualization: G.v.O., O.F. and S.J.C.; research design: O.F., G.v.O. and S.J.C.; experimentation: O.F.; data analysis: O.F.; Manuscript and figure preparation: O.F., S.H.S. and G.v.O.; Generation of lines: S.J.C and O.F.

---

**Figure S1. SA-induced period shortening is dose dependent.** Promoter activity of *CCA1* (A, E and I) and *TOC1* (C, G and J) was observed by measuring luminescence in CCA1pro:LUC and TOC1pro:LUC leaf disks, respectively. Leaf disks were treated with 100 µM SA (A and C), 10 µM SA (E and G), 10 mM SA (I and J) or mock (H_2_O). The mean period (B, D, F, H) was calculated from the 24 – 120 hr time window for each treatment. An unpaired t-test was performed between each mock and treated data set for *CCA1* and *TOC1*: *p <0.05, **p <0.01, ***p <0.001, ****p<0.0001, unpaired t-test. Error bars indicate mean ± SEM (n = 7). Data are from a single experiment representative of 3 independent repeats.

**Figure S2. Basal PR1 expression does not exhibit circadian variation.** Col-*0* seedlings were grown on MS plates under 12/12 light/dark conditions for 14 days before being released into constant light (LL). Every 3 hours, plates were sprayed with water. Relative expression of PR1 was measured 6 hours post treatment. The data points are technical replicates of quantitative PCR results combined from two biological replicates (orange versus blue data points; n = 3 each). The line represents mean values of all 6 replicates.

